# The Effect of Previously Encountered Sensory Information on Neural Representations of Predictability: Evidence from Human EEG

**DOI:** 10.1101/2025.05.27.656332

**Authors:** Kaho Magami, Roberta Bianco, Edward Hall, Marcus Pearce, Maria Chait

**Affiliations:** Ear Institute, University College London, London, WC1X 8EE, United Kingdom; Neuroscience of Perception & Action Lab, Italian Institute of Technology (IIT), 00161, Rome, Italy; School of Electronic Engineering and Computer Science, Queen Mary University of London, United Kingdom; Digital and Cognitive Musicology Lab, École Polytechnique Fédérale de Lausanne, Switzerland; Department of Clinical Medicine, Aarhus University, Denmark

**Author notes:** **Corresponding Author:** Kaho Magami, Ear Institute, University College London, 332 Gray’s Inn Road, London WC1X 8EE, UK; Maria Chait, Ear Institute, University College London, 332 Gray’s Inn Road, London WC1X 8EE, UK.

**Keywords:** Predictive Coding, Auditory scene analysis, Auditory memory, Regularity encoding, EEG

## Abstract

Accumulating evidence suggests that the brain continuously monitors the predictability of rapidly evolving sound sequences, even when they are not behaviorally relevant. An increasing body of empirical evidence links sustained tonic M/EEG activity to evidence accumulation and tracking the predictability, or inferred precision, of the auditory stimulus. However, it remains unclear whether, and how, this process depends on auditory contextual memory. In the present EEG study, we examined neural responses to sound sequences across two experiments, and compared them to predictions from ideal observer models with varying memory spans. Stimuli were sequences of 50 ms long tone-pips. In Experiment 1 (N=26; both sexes), a regularly repeating sequence of 10 tones (REG) transitioned directly to a different regular sequence (REGxREGy). In Experiment 2 (N=28; both sexes), the same regular sequence was repeated after an intervening random segment (REGxINTREGx). Results from Experiment 2 revealed that the inferred predictability of the resumed REGx pattern was influenced by the preceding INT tones, even several seconds after they ended, indicating that the brain retains contextual memory over time. In contrast, neural responses in Experiment 1 were best explained by models with minimal memory. This dissociation implies that the brain can dynamically adjust its strategy based on inferred environmental structure—resetting context when interruptions signal change, and preserving context when patterns are likely to resume.

## Introduction

The human brain is remarkably sensitive to the statistical regularities ubiquitously present in our surroundings (Arnal & Giraud, 2012; Bendixen, 2014; Bendixen et al., 2012; de Lange et al., 2018; Maheu et al., 2019; Press et al., 2020; Willmore & King, 2023; Winkler et al., 2009). A large body of research has demonstrated that observers can automatically acquire complex statistics from sensory inputs, including auditory, visual, and multimodal streams (Boubenec et al., 2017; Conway & Christiansen, 2005; Demarchi et al., 2019; Fiser & Aslin, 2001; Garrido et al., 2013; Horváth et al., 2001; Saffran et al., 1999; Stefanics et al., 2014; Turk-Browne et al., 2009; Wacongne et al., 2011). This computational ability is critical for generating predictions about the environment (Bendixen, 2014; Bendixen et al., 2012; de Lange et al., 2018; Friston, 2005; Press et al., 2020; Winkler et al., 2009), which allows the brain to optimize behavior by efficient allocation of cognitive and neural resources, supporting adaptive responses to incoming events (Bendixen et al., 2012; Boubenec et al., 2017; Bouwkamp et al., 2025; Kok et al., 2012; Nobre et al., 2007; Southwell & Chait, 2018; Yon et al., 2018).

An increasingly well-supported observation is that the neural mechanisms underlying the tracking of auditory statistical regularities can be studied through analyses of M/EEG sustained activity. These neural responses systematically vary with the predictability of sequential inputs, providing a direct window into how the brain monitors and adapts to environmental statistics (Barascud et al., 2016; Herrmann et al., 2019, 2021; Herrmann & Johnsrude, 2018; Hu et al., 2024; Southwell & Chait, 2018; Zhao et al., 2025).

Experiments using rapidly evolving auditory sequences have progressively revealed how the auditory system processes and accumulates statistical information about the acoustic environment. In the standard paradigm (e.g., Barascud et al., 2016), participants passively listen to tone sequences that transition between regular (REG) frequency patterns and random (RND) patterns. These sequences elicit a sustained neural response that dynamically tracks the structure of the auditory input (Barascud et al., 2016; Hu et al., 2024; Southwell et al., 2017; Zhao et al., 2025). Specifically, the emergence of a REG pattern is associated with a gradual increase in sustained neural activity, which plateaus as the regularity becomes established—suggesting that the brain has stabilized a representation of the repeating structure. Notably, longer and more complex patterns result in slower and more moderate amplitude increases, indicating limitations in the brain’s ability to discover and maintain representations of higher-order statistical regularities. Upon transition from REG to RND, the sustained response drops sharply and then settles into a lower, stable level—interpreted as reflecting the low predictability of random sequences. Zhao et al. (2025) extended these findings to stochastic sequences consisting of RND patterns with different predictability. These rises and drops in the sustained response align with predictions from computational ideal observer models (Harrison et al., 2020; Pearce, 2005; Skerritt-Davis & Elhilali, 2018, 2021), which quantify information content (IC; how surprising a given tone is based on prior exposure) or precision (inferred reliability of the predictive distribution; Yon & Frith, 2021) providing support for the hypothesis that the sustained response represents a mechanism that tracks predictability within the unfolding signal. However, it remains unclear how the brain determines the context or reference frame, whether derived from immediate sensory input or retrieved from longer-term memory, against which this predictability is assessed.

Commonly used modelling measures such as information content (e.g. as used in Harrison et al., 2020; Pearce, 2005), or precision (e.g. as used in Zhao et al. 2025), quantify the expectedness of an event given a particular context of previously encountered events stored in memory. In theory—drawing from Bayesian change-point estimation models (Adams & MacKay, 2007; Fearnhead & Liu, 2007; Wilson et al., 2010)—ideal observers should dynamically evaluate the relevance of a given context and determine how much of it to incorporate when constructing predictive distributions (Glaze et al., 2015; Nassar et al., 2010; Skerritt-Davis & Elhilali, 2018, 2021; Wilson et al., 2013). However, whether and how the sustained response, as a proxy for predictability processing, depends on experienced auditory events remains unexplored. Addressing these questions is crucial for understanding how past experiences are leveraged to represent the predictability of a given event.

Bianco et al. (2025) showed that REG patterns were recognized more quickly by the brain (as indicated by the MEG sustained response) when they were re-introduced following a scene interruption than when initially presented. This indicates the presence of an automatic memory store that carries a representation of the pattern across the interruption. More broadly, this finding suggests that by manipulating the information encountered by listeners and measuring its effects on the sustained response, it is possible to gain insight into what information is being stored and the conditions under which it is utilized. Here, we ask: Will the sustained response to a regular pattern be influenced by a listener’s prior experience with past information? This is tested by comparing two situations that differ in the relevance of prior experience: one in which a regularity is learned and then replaced by a new one—the prior experience is no longer relevant (Experiment 1), and another in which a regularity is learned, interrupted by a random tone sequence whose length is varied systematically, and then resumed—the prior experience is relevant and could be carried over (Experiment 2).

## Experiment 1

This experiment examines changes in the EEG sustained response triggered by transitions between two distinct regular (REG) patterns—REGx to REGy – compared with a continuation of REGx (Figure 1A). A similar comparison was made in one of the experimental conditions reported by Bianco et al. (2025) using MEG. Here, we replicate that approach using EEG to justify the use of EEG in the extension reported in Experiment 2.

**Figure 1.**
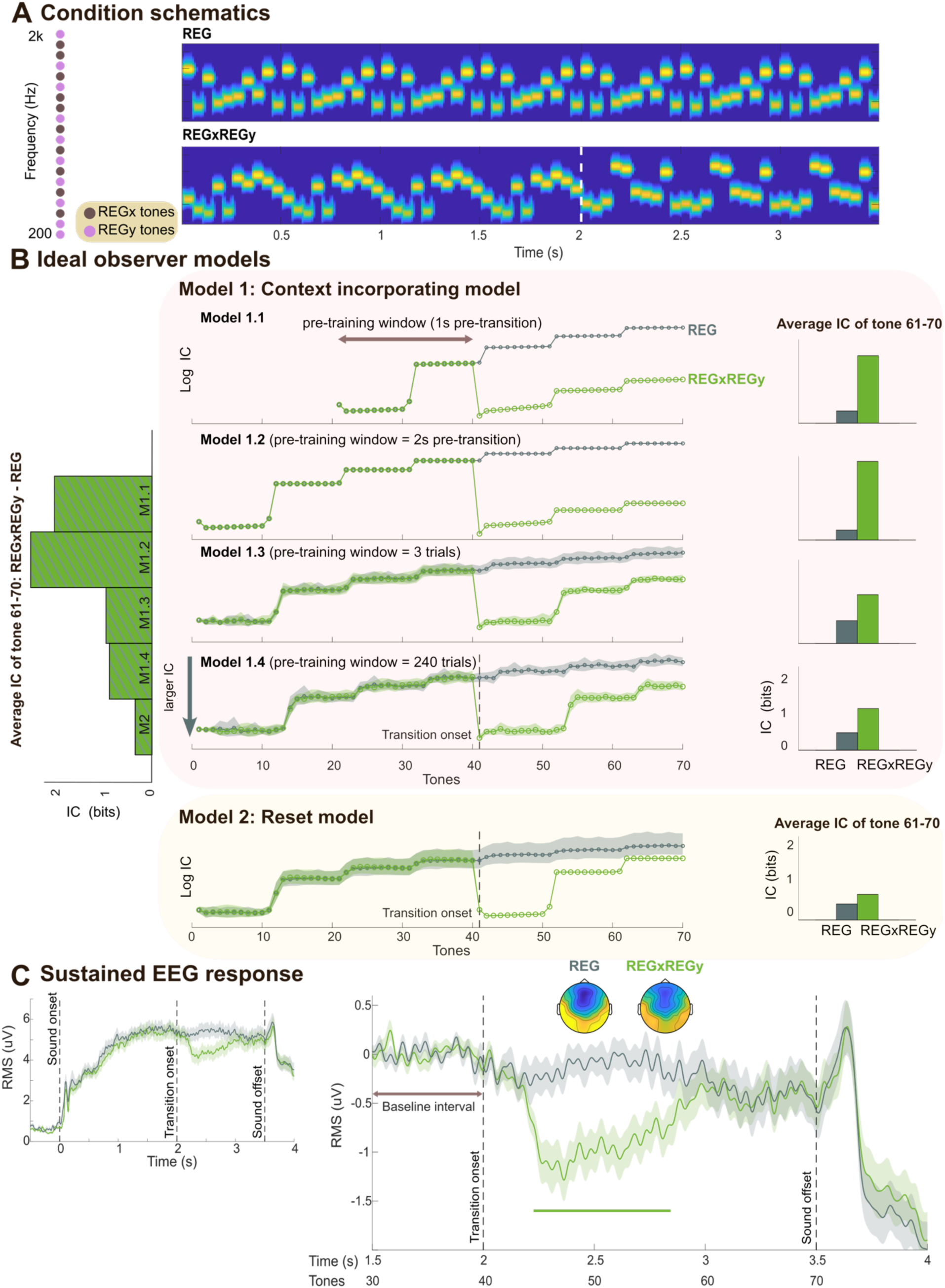
Experiment 1 stimuli, model simulations, and EEG results. **[A]** Left: Schematic illustration of the frequency selection method in Experiment 1, with each frequency represented as a circle. In this example, the brown frequencies were allocated to REGx and the pink to REGy. Right: Spectrograms depicting example stimuli for each condition. The dashed line marks the onset of REGy. **[B]** Model simulations. We implemented the model using the “new_ppm_simple” function from the ppm R package, available on GitHub (https://github.com/pmcharrison/ppm). All parameters were kept with their default settings, as described in the repository documentation. Middle: Information content (IC; log transformed) computed from variants of the IDyOM model, each incorporating different memory constraints. The y-axis is inverted (bottom = higher IC). The REG condition is shown in gray; REGxREGy condition is shown in green. For each condition, data are averaged over trials, with shaded areas representing twice the standard deviation (STDEV). Models vary by the duration of the “pre-training window”. Model 1.1 is pre-trained on 2 cycles of REGx (indicated by the brown arrow). Model 1.2 is pre-trained on 4 cycles of REGx. Model 1.3 is pre-trained over 3 trials. Model 1.4 is pre-trained over all 240 trials. Model 2 is reset upon pattern interruption, resulting in a pre-training window of length zero. All models estimate variable-order conditional probabilities for the next tone given the immediately preceding sequence of tones. The stimulus context over which the model learns representations of statistical structure that inform its conditional probabilistic predictions consists of the pre-training window and all tones experienced up to the time of prediction. The context varies between models, for example when predicting tone 50, the context is: Model 1.1, tones 20-49 of the current sequence; Model 1.2, tones 1-49 of the current sequence; Model 1.3, the three preceding trials plus tones 1-49 of the current sequence; Model 1.4, the 240 preceding trials plus tones 1-49 of the current sequence; Model 2, tones 41-49 of the current sequence. Right: Raw (non-log-transformed) IC values averaged over the last REGy cycle (tone 61 to 70; corresponding to 3 – 3.5 s). Left: IC differences (between REGxREGy and REG computed over tone 61-70) across all five models. **[C]** EEG data. Left: Group-averaged brain responses (RMS over 10 most responsive auditory channels; see Methods). Shading indicates twice the SEM (computed via bootstrap resampling, 1000 iterations). Data are baseline-corrected relative to the 0.5-second pre-onset window. Right: The same data but baseline-corrected using the 1.5–2 s pre-transition window. Significant differences (p<.01) between conditions are indicated by the horizontal bold line. Scalp topographies are based on activity averaged over the time window of significant differences (2.2–2.8 s relative to stimulus onset); the color ranges from −4 to 4 uV.

To inform our interpretation of the data, we use IDyOM, which implements a variable-order Markov model based on the Prediction by Partial Matching algorithm (Harrison et al., 2020; Pearce, 2005). The model has been extensively and successfully used to account for regularity processing in artificial sequences, such as those used in the current study (Barascud et al., 2016; Bianco et al., 2020, 2025; Harrison et al., 2020), as well as in more naturalistic musical settings (Cheung et al., 2019, 2023; Di Liberto et al., 2020; Kern et al., 2022; Quiroga-Martinez et al., 2021).

Starting with a null model, IDyOM learns incrementally based on the unfolding tone sequence and uses its learned model to generate a conditional probability distribution for each tone given the preceding tones. Figure 1B shows model predictions. To simulate the availability of different amounts of contextual information for REGy pattern detection, we varied the duration of the input sequence—referred to as the “pre-training window”—the model was trained on before the transition to REGy. In the simulations shown in Figure 1B, this window ranged from just a few seconds to 240 trials. Model predictions were always based on the full context available up to that point (i.e., all prior input; see figure legend). This approach allowed us to systematically manipulate the model’s memory content to examine how varying levels of prior information influence its output. The model quantifies the information content (IC) of each tone —reflecting the surprise elicited by that tone given the preceding context. We compare a *context incorporating model* that retains increasingly long spans of past input (from a few seconds to the entire experiment) with a *reset model* that clears its memory upon detecting a deviant tone—the first tone in REGy that violates expectations based on REGx.

All models show a gradual decrease in IC during REGx as the pattern is learned. This decrease occurs at different rates depending on the pre-training window; models with longer pre-training windows exhibit greater variability across trials (indicated by larger error bars) due to cumulative influences from prior tones. At the transition to REGy, all models show a sharp increase in IC, corresponding to the surprise elicited by an unpredictable tone. IC then remains high for a period before gradually reducing again, indicating that the new regularity (REGy) is being learned.

The duration of this learning period varies across models: models with a shorter pre-training window (e.g., Model 1.2) take longer to adapt (reflected by a slower decrease in IC), as existing memory content of REGx interferes with the encoding of the new pattern. Conversely, models with a longer pre-training window (e.g., Model 1.4) or where training is reset (e.g., Model 2) exhibit a more rapid decrease in IC, as the representation of REGy is less affected by prior memory of REGx.

This example also illustrates how the difference in IC between REGy and the non-changing REG control condition is modulated by the pre-training window. When the pre-training window is short, the difference in IC is consistently large, as the memory of REGx strongly influences the encoding of REGy. However, as the model’s pre-training window increases in length, this difference diminishes due to memory saturation: previously encountered patterns interfere with both REGx and REGy representations. In the *reset* model, the IC difference is also low, reflecting a complete lack of memory competition.

As discussed, the M/EEG sustained response is thought to reflect neural tracking of sequence predictability. If the brain represents sequence information similarly to the model, we would expect neural activity to mirror the dynamics of IC.

### Methods

#### Stimuli

The stimuli (Figure 1A) were 3500 ms long sequences composed of 50 ms tone pips (5 ms raised cosine ramps; 70 tone pips in total). REG sequences were generated by randomly selecting 10 frequencies from a pool of 20 logarithmically spaced values between 222 and 2000 Hz without replacement, and this sequence was cycled to create a regularly repeating pattern. For the REGxREGy sequence, two distinct REG sequences were generated: the first REG pattern (REGx) lasted 2 s, and the second (REGy) lasted 1.5 s. REGx was formed by randomly selecting 10 frequencies from the pool without replacement, and the remaining 10 frequencies were used to form REGy (Figure 1A). A unique sound sequence was generated for each trial and participant. The inter-stimulus interval (ISI) was jittered between 2.5 and 3 s.

#### Procedure

These data were collected as part of a separate study (Magami et al., in prep), that contained other stimuli (presented in a separate block).

Participants were seated in an acoustically shielded room (IAC triple-walled sound attenuating booth). They listened to auditory stimuli while engaging in a decoy visual task, presented on a computer screen located about 90 cm away. The visual task consisted of sequentially presented triplets of photographs of landscapes, and participants were instructed to press a key when the first and third images were the same (occurring in 40% of trials). Feedback regarding the number of hits, misses, and false alarms for the visual task was provided at the end of each block. The duration of the image presentation was jittered between 2 and 5 s, and images were cross faded to avoid abrupt visual transients. The timing of image presentation was not correlated with that of the auditory stimuli.

Overall, 120 sound stimuli were presented for each of the two sound conditions (REG, REGxREGy). These stimuli were presented randomly and arranged in 4 blocks. Sounds were presented diotically through headphones (3A Insert Earphone, 3M) via a Fireface UC sound card (RME) at a comfortable listening level (adjusted by each participant). Stimulus presentation was controlled with the Psychtoolbox package (Psychophysics Toolbox Version 3) in MATLAB (2019b The MathWorks, Inc.).

#### Recording, data processing, and statistical analysis

EEG signals were recorded using a Biosemi system (Biosemi Active Two AD-box ADC-17, Biosemi, Netherlands) from 64 electrodes at a sampling rate of 2048 Hz. Recording was restarted at the beginning of each block. For data analysis, the Fieldtrip (http://www.fieldtriptoolbox.org/) toolbox for MATLAB (2018a, MathWorks) was used.

The recorded data were down-sampled to 256 Hz, low-pass filtered at 30 Hz (two-pass, Butterworth, 5th-order) and detrended by a 1^st^-order polynomial. The data were divided into epochs of 6 s, from 1 s pre-stimulus onset to 1.5 s post-stimulus offset. The epochs were then baseline-corrected relative to the pre-onset interval (−0.5 s to 0 s relative to the sound onset). Outlier epochs and channels were removed by visual inspection, resulting in the removal of an average of 4.24 % of epochs and 0.9 channels per participant. De-noising source separation (DSS; De Cheveigné & Parra, 2014; De Cheveigné & Simon, 2008) analysis was then applied to each subject’s data across all conditions to maximize reproducibility across trials (over the interval of 0 s to 4 s relative to sound onset). For each participant, the first three DSS components were retained and projected back into sensor space. Finally, the data were re-referenced to the average of all channels, and the averages over epochs for each channel, condition and subject were calculated.

To quantify the effects, we selected the most auditory-responsive channels: for each participant, the N1 component (negative event-related potential happening at around 100 ms post-stimulus onset) of the sound onset response was identified from the averaged data. At the peak of the N1, the 5 channels showing the most positive activity and the 5 channels showing the most negative activity were considered to best reflect the brain’s auditory-related activity. In the figures below, we quantify the instantaneous power of the brain response by computing the RMS (root mean square) across these channels, following a similar approach in other works (Barascud et al., 2016; Southwell et al., 2017; Zhao et al., 2025). The RMS reflects the instantaneous power of the brain response regardless of polarity. Field maps at relevant time points are also provided.

#### Statistical analysis

To statistically evaluate the effect of interruption, the differences between sound conditions were calculated for each participant. This difference was then subjected to bootstrap resampling (Efron & Tibshirani, 1994). The difference between conditions was considered significant if the proportion of bootstrap iterations falling above or below zero exceeded 99% (p<.01) for more than 8 adjacent samples (Barascud et al., 2016).

#### Participants

Twenty-eight paid participants participated in Experiment 1. All reported no history of hearing or neurological disorders. Two participants were excluded due to exceptionally noisy EEG data. Data from the remaining twenty-six participants (19 females; average age 24.81, ± 4.20) were used for analyses. All experimental procedures were approved by the research ethics committee of University College London, and written informed consent was obtained from each participant.

### Results and discussion

#### The EEG sustained response tracks regularity discovery and violation

The group averaged responses for the two conditions (REG, REGxREGy) are shown in Figure 1C. Overall, we successfully replicated the findings from Bianco et al. (2025) with EEG.

The brain response exhibited an N1 peak at around 100 ms post-onset, then increased its amplitude until it reached a plateau before the end of the 2^nd^ cycle of the REG sequence. This sustained response pattern aligns with previous literature and is thought to reflect a rapid, automatic process of regularity detection (Barascud et al., 2016; Herrmann et al., 2019, 2021; Herrmann & Johnsrude, 2018; Hu et al., 2024; Southwell et al., 2017). Following the emergence of the REGy pattern, the sustained response rapidly dropped in amplitude, persisted at a low level (whilst the new REG pattern was being discovered) and then returned to the pre-transition level. To analyze the difference in the post-transition responses between conditions, we baseline-corrected the data relative to the pre-transition window (1.5-2 s post-onset; Figure 1C, right). Bootstrap resampling revealed a significant difference between the amplitudes of REG (control) and REGxREGy, starting from 220 ms (∼5 tones) after the interruption, consistent with the timing shown in the MEG data from Bianco et al. (2025).

As noted previously (Barascud et al., 2016; Bianco et al., 2025), unlike during regularity discovery, the EEG response latency here diverges from model predictions, which show a spike in IC immediately following the first tone that violates the REG pattern. Several factors could account for this divergence. One possibility is that the delay reflects a circuit-related delay in encoding the violation of the REG pattern. Alternatively, it might reflect a “wait-and-see” period, during which the system accumulates information about the scene change before responding. Indeed, Bianco et al. (2025) demonstrated that this latency is not fixed but scales with sequence information content (tone-pip duration), challenging the idea of a simple refractory period.

Following the abrupt drop in the sustained response, levels remained low for a period before rising again. The difference between conditions disappeared at 800 ms post-interruption (16 tones), at which point the response to REGy returned to the levels of the no change control condition (REG). Overall, these patterns indicate that the EEG sustained response dynamically tracks the brain’s process of discovering predictability, detecting its violation, and then fully re-establishing a new regularity.

#### Reconciling differences between modeling and the EEG response

There are notable differences between the EEG responses and the model’s behavior. For instance, as previously noted, the model exhibits an immediate response to the transition from REGx to REGy, whereas brain responses show a delay of about five tones.

A key point of divergence lies in how the model handles multiple cycles of regularity. Even in the control condition (no change, REG), the model continues to refine its representation of REG with each successive cycle. In contrast, the EEG sustained response to REG plateaus after approximately two cycles, indicating that the brain’s representation stabilizes relatively quickly.

As a result, in the model, REGy never reaches the same representational strength as REG in the control condition, since REG continues to be refined indefinitely. However, EEG data show that the transition from REGx to REGy leads to a return to the same level of sustained activity observed for REG within about one second of REGy onset. This discrepancy suggests that aspects of evidence accumulation—or more generally, auditory processing—that shape brain responses are not fully captured by the model.

Despite these differences, the models most consistent with the EEG findings are those in which the post-transition difference between REG and REGy is minimal—that is, models in which REGx and REGy do not strongly compete in memory. Such models typically either have a long pre-training window (e.g., Model 1.4) or are reset at the point of transition (Model 2), enabling a rapid reinstatement of a REGx-like response to REGy.

To further refine this interpretation, in Experiment 2, we asked how prior context affects the ‘rediscovery’ of a previously experienced regularity. To address this, we examined responses to an identical REG pattern while systematically varying the immediately preceding context.

## Experiment 2

We investigated the EEG sustained response evoked by an ongoing regular (REG) pattern occasionally interrupted partway. We employed a stimulus set (Figure 2A) in which 25% of the trials consisted of a regularly repeating sequence of tones. In the remaining trials, the regular pattern was interrupted by the insertion of 1, 3, or 5 novel tones (referred to as conditions INT1, INT3, and INT5, respectively) after which the original REG pattern resumed. We asked how this interruption would affect the representation of REG, with a specific focus on the speed at which the regularity was re-discovered and the post-interruption sustained response.

**Figure 2.**
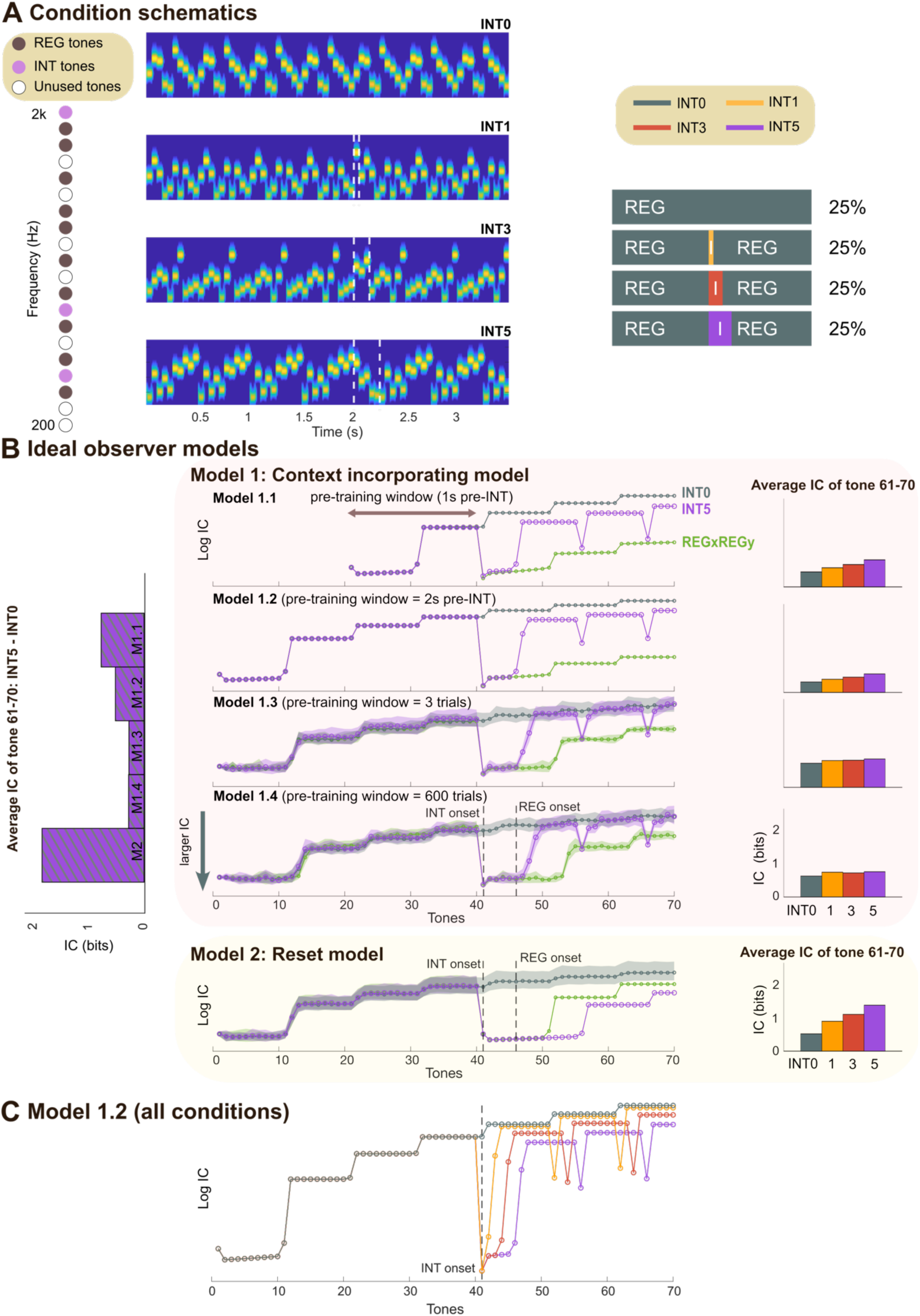
Experiment 2 stimuli and model simulations. **[A]** Left: Schematic of the frequency selection method in Experiment 2, with each frequency represented as a circle. The brown circles represent frequencies randomly chosen for REG; the pink circles represent tones chosen for the interruption tones (INT3 here). White circles denote unused frequencies in this trial. Middle: Spectrograms showing example stimuli for each condition. The white dashed box highlights the INT tones. The REG sequences before and after the INT tone follow an identical pattern. Right: Design schematics illustrating the stimulus sequences for each condition. ‘I’ indicates the interruption tones. **[B]** Model simulations. Middle: IC values (log-transformed) computed from variations of the IDyOM model, each with different memory constraints (as detailed in Figure 1), plotted for the INT0 and INT5 conditions from Experiment 2, and REGxREGy condition from Experiment 1. For each condition, data are averaged over trials, with shaded areas representing twice the standard deviation (STDEV). The y-axis is inverted (bottom = higher IC). Right: Raw (non-log-transformed) IC values averaged over the last REG cycle (tone 61 to 70; corresponding to 3 – 3.5 s). Left: IC differences (between INT5 and INT0 computed over tone 61-70) across all five models. **[C]** IC values for all four conditions, computed using Model 1.2.

As in Experiment 1, we constrained our hypothesis using IDyOM (Figure 2B). In this case, the stimuli consisted of a continuous REG pattern interspersed with occasional deviant tones. As a result, the IC differences between conditions are markedly smaller than those observed in Experiment 1 (Figure 1B). Nevertheless, the overall dynamics are consistent with those reported previously.

Importantly, the post-interruption behavior of the *context incorporating* model reveals two key phenomena: (1) Despite the post-interruption sequences being structurally identical across the INT conditions, IC levels remain distinct between them (Figure 2C). This occurs because the model incorporates the interruption tones into its predictive framework, increasing baseline uncertainty. (2) The model exhibits “phantom” IC spikes, reflecting an expectation for the interruption to recur. This behavior arises because the model lacks the capacity to infer higher-order rules, such as the one-time occurrence of interruptions and the guaranteed resumption of the pre-interruption pattern. Overall, the model’s behavior is dictated by its perfect memory of all prior experiences, with every past observation—regardless of its present relevance—being weighted equally. This includes the singleton interruptions, which continue to influence the model’s present IC estimates. This pattern is largely preserved across models with different pre-training window lengths, though models with longer pre-training show less pronounced differences in post-interruption IC (due to memory saturation; as discussed in Experiment 1, above).

For a model whose memory is reset at the interruption (Model 2), IC differences between conditions are also present because the ‘post-interruption world’ contains different numbers of unique elements for each interruption condition. The interruption tones are incorporated into model predictions, thereby decreasing baseline predictability. This model does not display phantom spikes, because its memory does not contain the previously experienced REG and its transition to the interruption tones.

Another difference between the models concerns the speed at which the REG pattern is re-discovered, reflected in the timing of the decrease in IC following the interruption. The *context-incorporating* models exhibit a rapid re-discovery of REG, whereas in the *reset* model, this process is slower due to the unavailability of pre-interruption memory.

Building on these insights, we examine whether passively listening participants exposed to these sequences will mirror model behavior. Specifically, we investigate human susceptibility to past interruptions, and whether these transient disruptions affect subsequent representations of regularity in a similar manner to modelling.

### Methods

#### Stimuli

The stimuli were 3500 ms long sequences of 50 ms tone pips (5 ms raised cosine ramps; 70 tone pips in total). Tone frequencies were drawn from a pool of 20 logarithmically spaced values between 222 and 2000 Hz. Each stimulus comprised a sequence of regularly repeating tones (REG), generated in the same manner as in Experiment 1 (Figure 2A). In 25% of trials, the REG pattern continued with no interruption (INT0). In the remaining trials, an interruption in the form of 1, 3, or 5 new tones was introduced at 2000 ms post-onset (following 4 cycles of REG). These conditions will be referred to as INT1, INT3 and INT5, respectively. The frequencies of INT tones were randomly selected without replacement from the pool of remaining frequencies not used to form the REG sequence. Following INT, the original REG pattern was re-started. The duration of this remaining portion varied across conditions (1500 ms, 1450 ms, 1350 ms, and 1250 ms for INT0, 1, 3, and 5 conditions, respectively), ensuring that the overall tone number remained fixed at 70 tones. The ISI was jittered between 2.5-3 s. A unique sound sequence was generated for each trial and participant.

#### Procedure

General procedures were identical to those in Experiment 1. Overall, 600 sound stimuli were presented (150 stimuli per condition; in random order). The session was divided into 5 blocks, each approximately 10 min long. Participants were allowed a short rest between blocks.

#### Recording and data processing

General protocols were identical to those described in Experiment 1. On average, 1.47% of epochs were removed as outliers, along with 0.5 channels per participant. For the detailed comparisons of RMS values between conditions, we applied two different baseline correction time windows to the output RMS. For the comparison of the post-interruption neural response, we applied baseline correction at the time window before the interruption onset (1.5 s to 2 s post-onset). To compare the timing where the neural response after interruption tones stabilizes, we applied baseline correction at a different time window (3 s to 3.3 s post-onset). Additionally, post-interruption neural responses were compared across INT conditions (INT1, INT3, and INT5) by subtracting the control condition (INT0) from each INT condition, followed by baseline correction in the 1.5–2 s post-onset window.

To uncover activity potentially masked by the slow DC changes, the same analysis was performed on high-pass filtered data at 2 Hz (two-pass, Butterworth, 4th-order) with baseline correction applied just before the onset of the interruption (1.8 s to 2 s post-onset). DSS was applied to the data around the interruption tone (1.5 s to 4 s post-onset), and 2 components of the DSS outputs were retained for the data representation. As we were especially interested in the mismatch negativity (MMN)-like response to the pattern interruption, we selected the electrodes best reflecting the MMN response. To do this, we averaged the data across all conditions across all participants and selected the 10 electrodes with the most negative activation at the typical MMN response time (150 ms to 200 ms post-interruption-onset).

To examine the possible presence of the “phantom” interruption peaks, we analyzed the high-pass filtered data at 2 Hz by applying DSS separately to each experimental condition (2 s to 4 s post-onset) and extracted 2 components for the data representation. Analyzing each condition separately was necessary because the “phantom” peaks occur at a different latency in each condition (tone 52 and 62 in INT1, 54 and 64 in INT3, and 56 and 66 in INT5; Figure 2B, C). The output data were then segmented into 600 ms epochs (from 400 ms before the model-based peak timing to 200 ms after the peak timing) to limit the analysis at around the model-inferred peak locations. The initial 200 ms of each epoch was used for baseline correction, and the responses from 10 channels (same as those used in the RMS calculation) were averaged.

#### Statistical analysis

To statistically evaluate the effect of interruption, the differences between sound conditions (INT0, INT1, INT3, and INT5) were calculated for each participant. This difference was then subjected to bootstrap resampling (Efron & Tibshirani, 1994). The difference between conditions was considered significant if the proportion of bootstrap iterations falling above or below zero exceeded 99% (p<.01) for more than 8 adjacent samples (Barascud et al., 2016).

For the bootstrap analysis on data baseline-corrected to 3–3.3 s post-onset, we defined the final significance point as the moment when the neural response stabilized. To assess whether this timing differed across conditions, we repeated the bootstrap analysis for each condition pair (INT0 vs. INT1, INT0 vs. INT3, and INT0 vs. INT5). Specifically, we performed 1000 iterations of bootstrap resampling per pair and identified the last significant data point within the interval from INT offset to 3 s (the onset of the baseline correction window) in each iteration.

#### Participants

Thirty paid participants participated in Experiment 2. All reported no history of hearing or neurological disorders. Two participants were excluded due to exceptionally noisy EEG data. Data from the remaining twenty-eight participants (22 females; average age 23.4, ± 3.41) were used for analyses. All experimental procedures were approved by the research ethics committee of University College London, and written informed consent was obtained from each participant.

### Results and discussion

#### All interruption conditions elicit early MMN-like responses

Sensitivity to the interruption was evaluated by analyzing the response at the transition. To isolate the MMN-like response which we expected to be evoked by the INT (deviant) tones (Figure 3), the EEG data were high-pass filtered at 2 Hz and averaged across trials for each condition (as detailed in the Methods section). Indeed, the response is not visible in the non-high-pass-filtered data; see Figure 4. Bootstrap resampling revealed significant deflection in the INT1, INT3, and INT5 conditions relative to the INT0 condition (Figure 3), with latencies emerging between 70-100 ms post interruption onset. Notably, this latency and the corresponding topography (Figure 3; bottom) are consistent with those commonly associated with the MMN response (Winkler, 2007). Overall, this suggests that the interruption was similarly detected by the brain in all conditions.

**Figure 3:**
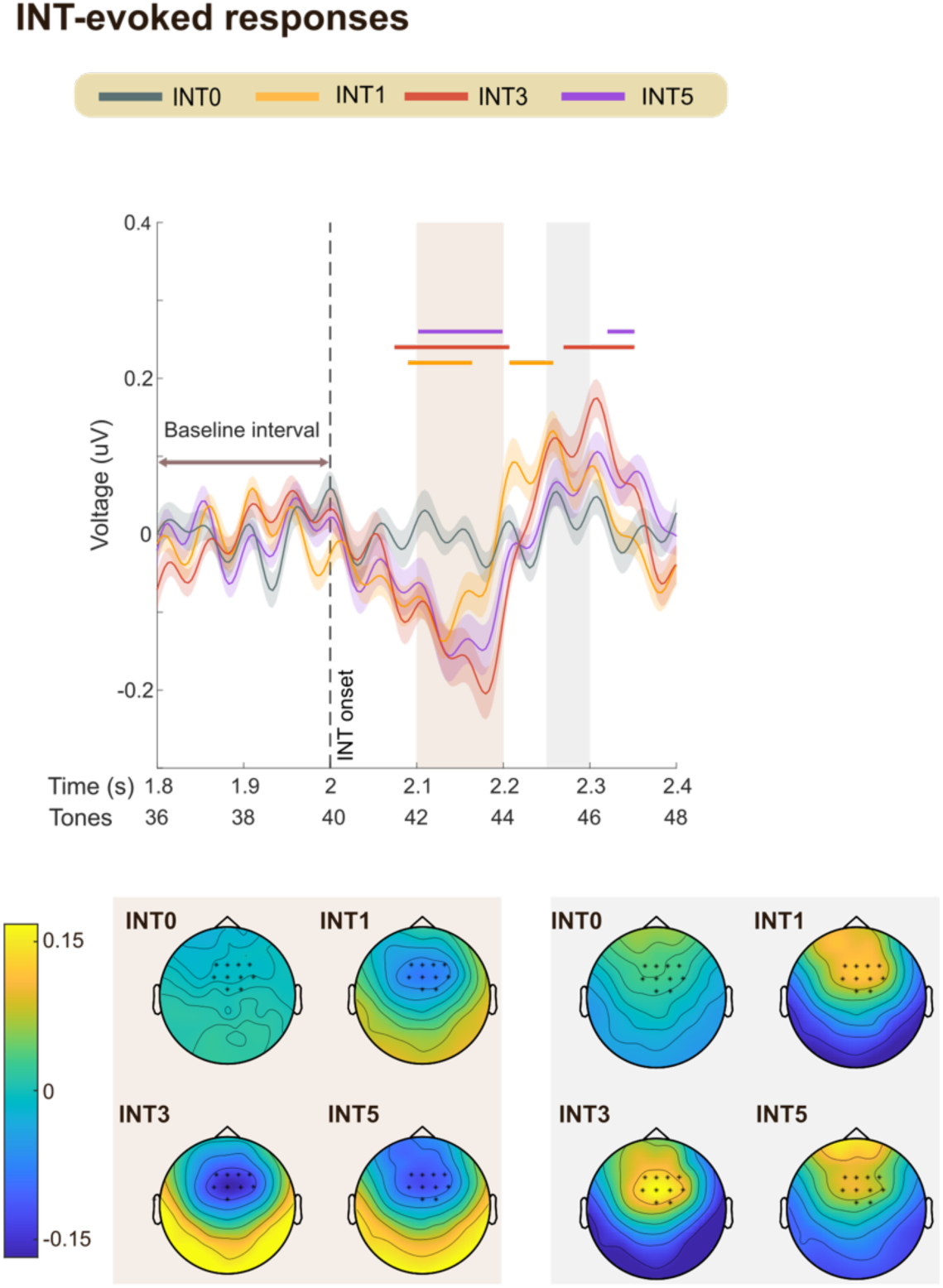
Experiment 2: INT evoked deviance response. High-pass filtered mean EEG data, averaged over 10 channels (indicated in the scalp topographies). Shaded areas represent twice the SEM. Significant differences (p<.01) between INT0 and interruption conditions (INT1, INT3, INT5) are indicated by the horizontal lines above the EEG traces. Scalp topographies, calculated for two time windows (2.1-2.2s and 2.25-2.3s), are shown at the bottom.

**Figure 4.**
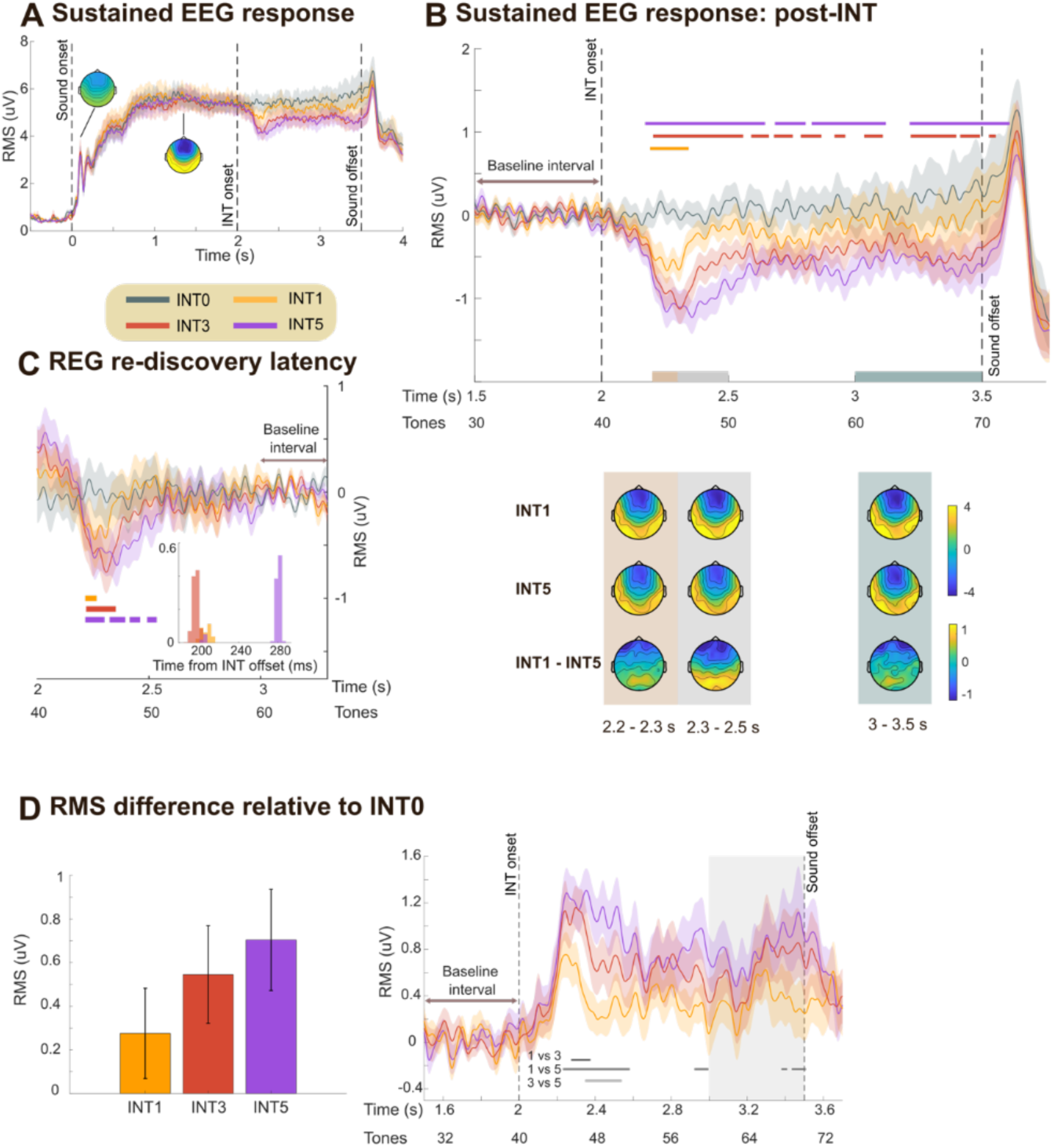
Experiment 2: sustained response dynamics. **[A]** Group-averaged RMS of brain responses. Shaded areas represent twice the SEM. Data are baseline-corrected to the −0.5–0 s pre-onset window. Scalp topographies illustrate two response phases: N1 component (80–150 ms post-sound onset) and the sustained response (1–2 s post-sound onset); the color ranges from −4 to 4 uV. **[B]** Same data as in [A] but baseline-corrected to the pre-interruption window (1.5–2 s). Significant differences (p<.01) between INT0 and INT1, INT3, INT5 are indicated by bold horizontal lines above the EEG traces. Scalp topographies are provided for three time windows: 2.2-2.3 s, 2.3-2.5 s, and 3-3.5 s relative to sound onset. **[C]** Same data as in [A], baseline-corrected to 3–3.3 s. Significant differences (p<.01) between INT0 and INT1, INT3, INT5 are indicated by bold lines below the EEG traces. The histogram (inset) shows the latencies associated with REG re-discovery. The results demonstrate delayed rediscovery of REG in the INT5 condition. **[D]** Right: EEG data (RMS) for each of the interruption conditions after subtracting the INT0 condition, baseline-corrected within the pre-transition window (1.5–2 s). Grey lines beneath the traces mark significant differences (p<.01) between INT conditions (INT1 vs. INT3, INT1 vs. INT5, INT3 vs. INT5). Left: Same data averaged over the 3–3.5 s time window (gray shading). Error bars represent SEM.

**Figure 5.**
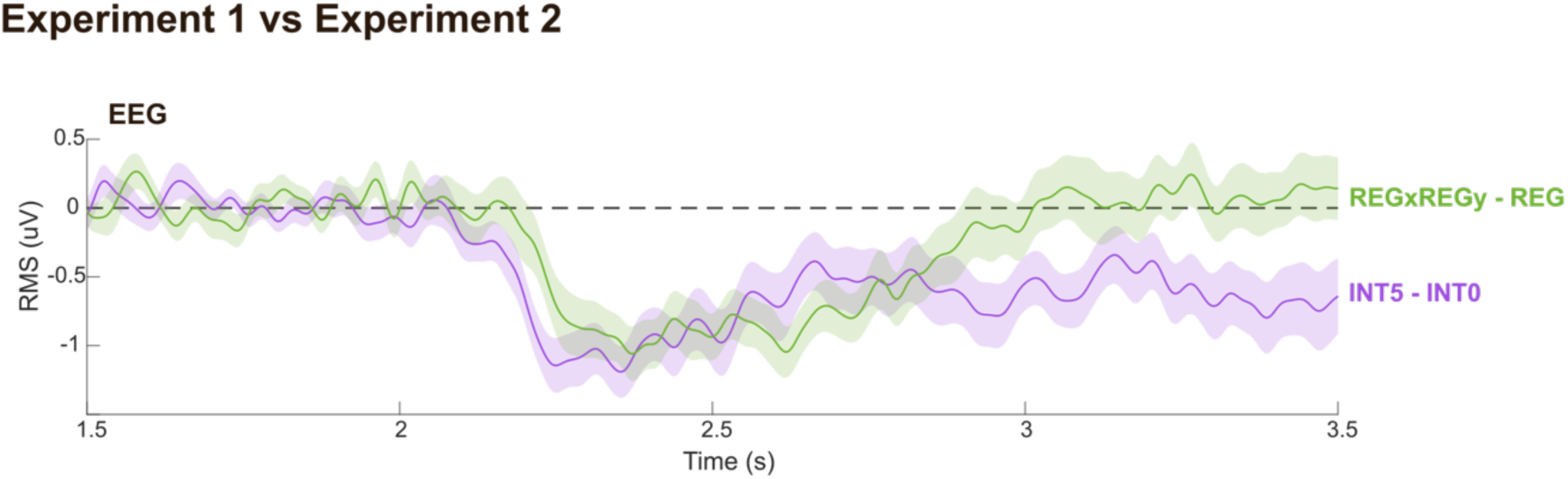
Comparing EEG across Experiments. Group-averaged RMS of brain responses. The difference between REGxREGy and REG (Experiment 1) is shown in green. The difference between INT5 and INT0 (Experiment 2) is shown in purple. Data are baseline-corrected using the pre-transition window (1.5–2 s). Shaded areas indicate ±2 SEM.

#### The EEG sustained response tracks the dynamics of sequence IC

For each participant and condition, the RMS over 10 selected channels (detailed in the Methods section) was calculated at each time point on the non-high-pass filtered data. The group averaged RMS amplitude for the four conditions (INT0, INT1, INT3, and INT5) is shown in Figure 4A. The general trajectory mirrored the pattern observed in Experiment 1: the sustained response increased and plateaued as the brain adapted to the REG pattern, dropped in amplitude following the INT tones, and then recovered (but not fully to baseline) as the brain re-engaged with the REG pattern. This trajectory aligns with the information content patterns predicted by the IDyOM models.

Following the REG interruption, the sustained response dropped rapidly. To analyze differences in post-interruption responses across conditions, we baseline-corrected the data relative to the pre-interruption window (1.5–2 s post-onset; Figure 4B). Bootstrap resampling (see Methods) revealed a significant difference between the control condition (INT0) and the interruption conditions (INT1, INT3, and INT5), starting at 187 ms (∼4 tones) after the interruption onset. This finding is consistent with previous observations (Barascud et al., 2016; Bianco et al., 2025), including Experiment 1 in this study.

As noted previously, one possibility is that the delay reflects a fixed refractory period after the MMN-like response or some other circuit-related delay in encoding the violation of the REG pattern. Alternatively, it might reflect a “wait-and-see” period. If the 4-tone latency reflects a period of assessment—during which the system evaluates whether the REG violation is a spurious event or indicative of a consistent stimulus change—we might expect no interruption response in INT1 but a larger response in INT5. This was partially observed: while all interruption conditions exhibited a similar latency for the sustained response drop, the trough was deeper for INT3 and INT5 than for INT1 (Figure 4D)

#### The EEG sustained response indicates a memory trace for REG post interruption

To assess the time required to re-learn the REG pattern—reflected in the recovery of the sustained response—we baseline-corrected the data relative to the post-recovery window (3–3.3 s post-onset; see indicated in Figure 4C). Bootstrap resampling (see Methods) identified time points where responses to the interruption conditions (INT1, INT3, and INT5) remained significantly below the control (INT0) condition. We defined amplitude recovery as the latest time point where this significant difference was observed. This occurred approximately 216 ms (∼4 tones), 202 ms (∼4 tones), and 285 ms (∼5 tones) after the offset of the final interruption tone in the INT1, INT3, and INT5 conditions, respectively.

Under perfect memory conditions—as seen in the model dynamics—the timing of REG re-discovery following the interruption should be the same across all INT conditions, once the duration of the interruption is accounted for (i.e., subtracting 1, 3, or 5 tones, respectively; e.g., see Figure 2C). In contrast, our results show a longer latency following INT5 compared to INT1 and INT3. Bootstrap resampling confirmed a consistent difference between the INT1/INT3 and INT5 conditions (Figure 4C), suggesting that INT5 requires one additional tone to re-establish the REG pattern after REG is reintroduced. This may reflect neural memory constraints that limit the speed of pattern re-learning.

Critically, and notwithstanding the differences between conditions highlighted above, the observed recovery times were consistently shorter than a regularity cycle (i.e., <10 tones) and faster than model predictions for the discovery of a new REG pattern (i.e., 1 cycle + ∼5 tones, as observed in Experiment 1). This indicates that, despite the interruption, the brain retained a memory of the REG pattern, enabling faster re-discovery (see also Bianco et al., 2025).

#### Persistent post-interruption sustained response differences between conditions

In contrast to the results in Experiment 1, after the interruption, persistent differences in the sustained response between INT 1, 3, 5 and INT0 were observed; Figure 4B indicates that sustained responses did not return to the pre-interruption baseline in INT conditions. Given that the amplitude of the sustained response is hypothesized to reflect the brain’s representation of the predictability of unfolding sounds, this reduced amplitude suggests a decrease in inferred predictability with exposure to a greater number of INT tones, similar to that observed in modelling.

One salient feature in the model is the expectation of “phantom” interruption events at the onset of every regularity cycle following the pattern interruption (tone 52 and 62 in INT1, 54 and 64 in INT3, and 56 and 66 in INT5, as shown in Figure 2B and C). Our analysis (see Methods) did not yield consistent EEG parallels. We cannot rule-out the possibility that the noisy nature of EEG signals obscured them and that the fluctuations in the time domain (e.g. see Figure 4B) are a smeared manifestation of these peaks.

The general pattern of a speeded re-discovery of REG and a persistent lower sustained response in the INT conditions matches the predictions of the “context incorporating” family of models. This is because the *reset* model does not predict faster re-discovery of REG, and the *context incorporating* models maintain a memory of the INT tones that directly affect the representation of the REG sequence following its resumption.

The mean amplitude patterns across INT conditions (Figure 4D, left) were consistent with an effect of INT duration on the sustained response, although there was no statistically significant difference between the INT conditions (*F*(2,54) = 1.81, p = 0.17; repeated-measure ANOVA). The difference from the control (INT0) appeared graded, as reflected in the pattern of significance (horizontal lines) in Figure 4B. Direct condition comparisons revealed only a small effect between INT1 and INT5 (Figure 4D, right). This is perhaps not surprising given that the conditions only differed by the introduction of 2 tones. But overall, we can conclude that the pattern of EEG data is consistent with a model that maintains a long enough pre-training window to incorporate a memory of the preceding REG and the INT tones into the inferred predictability of the post-interruption REG.

Overall, these results indicate that the presence of interrupting tones affected the representation of REG even a second or more after the interruption had ended. This pattern aligns with the predictions of *context incorporating* models (Model 1) which suggest that memory of the INT tones influences the IC of the REG pattern in a manner reflected in the EEG data.

## General Discussion

Our analysis focused on the dynamics of the EEG sustained response. Accumulating evidence suggests that it reflects the process of predictability tracking in statistically structured sequences (Barascud et al., 2016; Bianco et al., 2025; Hu et al., 2024; Zhao et al., 2025), supported by the coordinated processing of information across a distributed neural network. Source localization of the MEG sustained response (Barascud et al., 2016; Bianco et al., 2025; Hu et al., 2024) implicates a distributed network involving the auditory cortex (AC), hippocampus (HC), and inferior frontal gyrus (IFG) in representing REG patterns. This activity fluctuates dynamically, decreasing during REG interruptions and reinstating upon the discovery of a new REG pattern. These fluctuations likely reflect the disruption of top-down connectivity when an existing model is deemed no longer relevant and the strengthening of top-down connectivity when predictive models are available.

We investigated whether and how the passive-listening brain utilizes past experiences to represent ongoing sound sequences by recording EEG sustained responses in two situations: one in which a REG sequence is replaced by a different REG sequence (Experiment 1) — and another in which a REG sequence is occasionally disrupted by a varying number of new tones (Experiment 2).

### Sustained responses to REG patterns are affected by brief interruptions

Experiment 2 revealed that sustained responses to the post-interruption REG patterns were affected by the INT tones. This finding suggests that the brain represents the post-INT REG sequence using past information, including the history of INT tones.

Prior research shows that the brain incorporates long-term sensory history when processing sequences (Maheu et al., 2019; Rubin et al., 2016; Ulanovsky et al., 2004; see also Demarchi et al., 2019; Fritsche et al., 2022), though estimates of this duration vary depending on the specifics of the paradigm used. For instance, Rubin et al.(2016) found that auditory cortex neurons in anesthetized cats best fit prediction models accounting for more than ten previous tones (∼9 seconds). Similarly, Benjamin et al. (2024) showed that tone information remains decodable from MEG responses for approximately eight successive items (2 seconds) during passive listening. Skerritt-Davis and Elhilali (2018) revealed that memory span, estimated by fitting a Bayesian perceptual model to behavioral data, correlated with performance, extending up to the full duration of each sequence (60 tones; ∼19 seconds). Zhao et al. (2025) applied a similar model to random tone-pip sequences and found that most listeners based their judgments on a context of 20–40 tones (∼1-2 seconds).

Here, we show that even brief contextual perturbations—such as a single interrupting tone—can alter the brain’s representation of an ongoing pattern. Given the link between the sustained response and perceived predictability, the reduced sustained response amplitude following interrupting (INT) tones indicates that the inferred predictability of the REG pattern was diminished after the interruption, despite no change in the stimulus itself.

This phenomenon—where transient surprise alters neural responses to an otherwise unchanged stimulus—has been observed across multiple research domains. In post-traumatic stress disorder (PTSD), for example, neural and physiological responses to a stimulus can change after a surprising or stressful event coincides with it, even if the stimulus itself remains the same (Kaczkurkin et al., 2017; Nutt & Malizia, 2004; Sartory et al., 2013; Wessa & Flor, 2007). Similarly, in perceptual decision-making, when participants predict an image’s location or value based on previous patterns, a surprising rule deviation can significantly alter their representation of the stimulus and its environment (Kao et al., 2020; McGuire et al., 2014; Nassar et al., 2010, 2012). This consistency across different psychological domains suggests a fundamental heuristic employed by the brain to track environmental changes.

### Sustained response dynamics reflect memory of INT and pre-interruption REG

As discussed above, the sustained response modulations are consistent with a memory trace for the interruption, which causes the sustained response to settle below the level observed in the control condition (INT0). As seen also in the model, the same memory effects also influence the speed of REG re-discovery after INT. Specifically, re-discovery occurs more rapidly than the initial discovery of a new regularity (see also Bianco et al., 2025). In all cases, the sustained response begins to rise before a full cycle of the REG pattern has elapsed. Thus, both the modulation of the sustained response and the accelerated re-discovery of REG following INT reflect the system’s use of prior information, aligning with the notion of a “context-incorporating” memory process, even within a simplified model framework.

Interestingly, the response dynamics also suggest memory decay. According to the model, under perfect memory conditions, the latency of REG re-discovery should be identical across INT conditions once the duration of the interruption is accounted for. However, this is not what we observe in the EEG data: following INT5, the re-discovery is delayed by approximately one tone (50 ms) compared to shorter interruptions. This delay cannot be explained by increased memory interference, as each INT condition uses a distinct set of tones, eliminating overlap. Instead, the delay points to a reduction in memory duration—i.e., decay—rather than a loss of memory content. This finding is significant because it demonstrates that memory decay can be detected within this paradigm, even during passive listening.

### Distinct sustained response patterns in Experiment 1 and Experiment 2 suggest listeners can use or ignore context depending on its relevance

Experiment 1 yielded a different pattern of results to Experiment 2, despite the use of similar sound stimuli and the same analysis protocol. While acknowledging that Experiments 1 and 2 were conducted separately and differ in several respects—making direct comparisons necessarily speculative—this divergence suggests that the passive-listening brain may employ a flexible context integration strategy.

In Experiment 1, the sustained response indicated that the REGy representation remained unaffected by the REGx context, as reflected in the full recovery of REGy amplitude to the REGx level (Figure 1). This pattern aligns with models where the REGx memory was either diluted by other contextual memories or erased entirely. In our simple modelling world, the results of Experiment 1 are either consistent with the *reset* model (Model 2) or a *context incorporating* model that learns from a relatively long prior context (e.g. Model 1.4).

In contrast, as discussed above, the pattern of results in Experiment 2 is not consistent with a *reset* model, but rather with models maintaining a memory of the preceding trials. Notably, as illustrated in Figure 2, *context incorporating* models predict a greater IC deviation from the control condition (REG, INT0) in the REGxREGy condition (Experiment 1) than in the INT5 condition (Experiment 2). However, the EEG data reveal the opposite pattern—the INT5 condition shows a larger deviation from the control. Therefore, to reconcile both experiments, a parsimonious conclusion is that different strategies are used by the brain in the two experiments: a “memory reset” strategy for Experiment 1 and a “memory incorporating” strategy for Experiment 2.

One possible explanation for the existence of different strategies in the two experiments lies in the distinct differences in stimulus set, and hence listeners’ belief about the environment imposed in the two experiments. In Experiment 1, a violation of REGx (the first tone violating the REGx pattern) was always associated with a transition to a new pattern (REGy), meaning that REGx was not relevant, and its representation could be discarded to facilitate the learning of REGy. In contrast, in Experiment 2, the regular pattern consistently re-emerged shortly after an interruption, reinforcing the expectation that the pre-interruption REG sequence remained relevant. Therefore, in Experiment 1, participants may have automatically adapted by discarding the REGx representation, similar to the behavior of the *reset* model, which erases prior context when making new predictions. Conversely, in Experiment 2, participants may have learned that the pre-interruption REG pattern remained relevant, leading them to preserve its memory even after the interruption. This difference suggests the brain’s ability to flexibly adjust its memory integration strategies based on the statistical structure of the auditory environment.

Importantly, a similar effect was also observed in Bianco et al. (2025; Figure 6 reproduces their findings). In that study, the authors investigated two experimental auditory environments. One, labelled ‘ENVnovel’, consisted of sequences that always transitioned to a new pattern (REGxREGy, as in the current study; REGxRND; and a REGx control). The other context, ‘ENVresume’, presented in a separate set of blocks, included REGxRND and REGx sequences but also, crucially, a condition in which the original REGx pattern resumed after an interruption by 10 random tones (REGx-RND-REGx; Unlike in the present Experiment 2, the length of the interruption was not varied). Thus, while in ENVnovel it was possible to “discover” that once REGx was disrupted, the associated predictive model was no longer relevant (since REGx would not reappear), this was not the case in ENVresume, where REGx was reintroduced 30% of the time. The results of that experiment revealed a pattern consistent with the findings reported here: the sustained response in the REGx-RND-REGx condition was consistently lower than in its control counterpart. This suggests that the brain retains information about past regularities in memory even when they are not guaranteed to reoccur.

**Figure 6.**
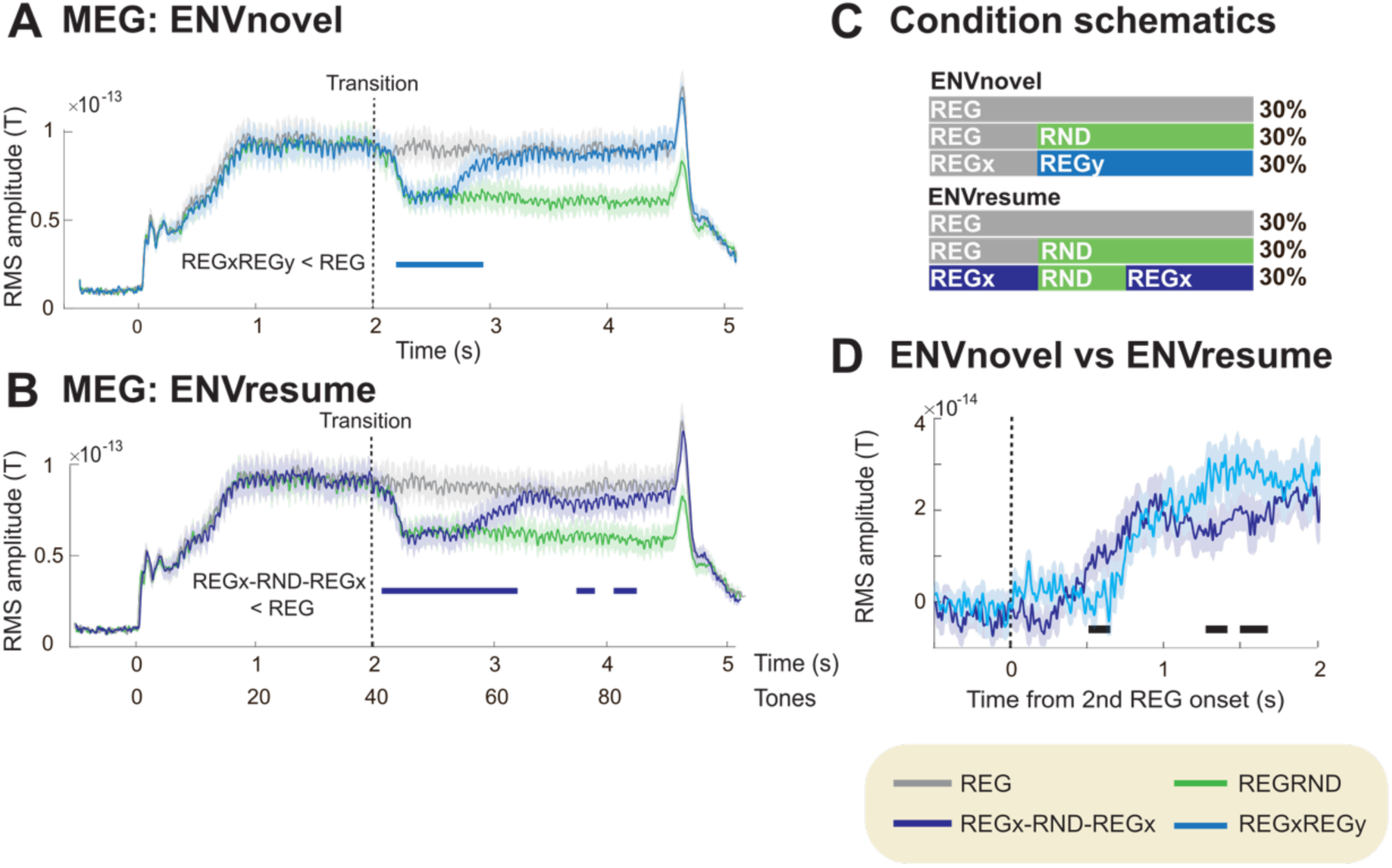
Results from Bianco et al. (2025) reveal results consistent with Experiment 2 here. The study presented stimuli in two contexts. In ‘ENVnovel’, transitions were always to a new pattern (REGxREGy, as in the current study; REGRND; and a REG control). ‘ENVresume’, presented in a separate set of blocks, included REGRND and REG sequences but also, crucially, a condition in which the original REGx pattern resumed after an interruption by 10 random tones (REGx-RND-REGx). **[A]** Group-average MEG brain responses (RMS across “auditory” channels; see more details in Bianco et al. (2025)) from the ENVnovel block. Data are baseline-corrected to the −0.5–0 s pre-onset window. Significant differences are indicated by the bold line below the MEG traces. These results are consistent with those in Experiment 1 here. **[B]** Data from the ENVresume block, demonstrating a persistent difference between REG and REGx-RND-REGx conditions following the resumption of REGx. **[C]** Design schematics illustrating the stimulus sequences for each condition. **[D]** Direct comparison of REGxREGy from ENVnovel and REGx-RND-REGx from ENVresume. REGRND condition data are subtracted from each condition of interest. Significant differences between conditions are indicated by bold lines below the traces. The results demonstrate a reduced sustained response in ENVresume relative to ENVnovel, consistent with the observations in Experiment 1 and 2 here.

This flexibility in adjusting the duration of reference memory is considered a crucial feature of the brain, allowing it to maintain an accurate representation of a dynamically changing environment (Bland & Schaefer, 2012; Glaze et al., 2015; O’Reilly, 2013; Yu & Dayan, 2005). Such adjustments occur in response to environmental state changes or change points—moments when past observations become unreliable for predicting future events. When a change point occurs, minimizing the influence of past memory and prioritizing new evidence accumulation enables a rapid adaptation to the new environment (Glaze et al., 2015; Nassar et al., 2010; O’Reilly, 2013; Skerritt-Davis & Elhilali, 2018, 2021). Empirical studies, mostly conducted using tasks involving slow decision-making and active attention allocation, suggest that humans can flexibly adjust their change point assumptions based on volatility estimates (Behrens et al., 2007; Glaze et al., 2015, 2018; Nassar et al., 2010). The present results suggest that similar heuristics might be operating on a faster time scale associated with sensory processing. Further controlled studies are needed to clarify these effects. For example, comparing results from conditions (REGx-INT-REGy vs REGx-INT-REGx) presented in separate blocks versus intermixed within the same block could help determine whether strategy adjustments occur on a trial-by-trial basis, at the block level, or require prolonged exposure throughout the experiment.

Additionally – it is important to stress that we use simple model comparisons focusing on varying the length of the pre-training window (consisting of counts of occurring n-grams). However, we did not account for other model dynamics or potential parameter variations. Moreover, while we used the IDyOM model due to its success in predicting human sequential processing (Barascud et al., 2016; Cheung et al., 2019; Di Liberto et al., 2020; Kern et al., 2022; Quiroga-Martinez et al., 2021), this model does not account for complex cognitive constraints, such as dynamic memory limitations and low-level auditory sensitivity. Further exploration of various models and model parameters is critical for our understanding of how the brain dynamically tracks the ongoing sequences under dynamic environments.

## Conflicts of interest

The authors declared no conflicts of interest concerning the research, authorship, and/or publication of this article.

## Acknowledgments

This work was supported by a BBSRC project grant to MC. KM is funded by Japan Student Services Organization. The funders had no role in study design, data collection, and analysis, decision to publish, or preparation of the manuscript.

## Data sharing

The data reported in this manuscript alongside related information will be available on OSF upon publication.

## Notes

### Competing Interest Statement

The authors have declared no competing interest.

